# Horizontal Gene Transfer Constrains the Timing of Methanogen Evolution

**DOI:** 10.1101/129494

**Authors:** Joanna M. Wolfe, Gregory P. Fournier

## Abstract

Microbial methanogenesis may have been a major component of Earth’s carbon cycle during the Archaean Eon, generating a methane greenhouse that increased global temperatures enough for a liquid hydrosphere, despite the sun’s lower luminosity at the time. Evaluation of potential solutions to the “faint young sun” hypothesis by determining the age of microbial methanogenesis was limited by ambiguous geochemical evidence, and the absence of a diagnostic fossil record. To overcome these challenges, we utilize a temporal constraint: a horizontal gene transfer (HGT) event from within archaeal methanogens to the ancestor of Cyanobacteria, one of the few microbial clades with recognized crown group fossils. Results of molecular clock analyses calibrated by this HGT-propagated constraint show methanogens diverging within Euryarchaeota no later than 3.51 Ga, with methanogenesis itself likely evolving earlier. This timing provides independent support for scenarios wherein microbial methane production was important in maintaining temperatures on the early Earth.

## Introduction

Methane is a greenhouse gas implicated in current and past climate change. Accumulation of atmospheric methane during the Archaean Eon has been proposed as one solution to the “faint young sun paradox”, contributing to increased global temperatures enough to maintain a liquid hydrosphere despite the lower luminosity of the sun at the time^1,2^. While microbial methanogenesis is generally assumed to be an extremely ancient pathway due to its phylogenetic distribution across much of Euryarchaeota^3^, there is only limited geochemical evidence for microbial methane production in the Archaean, in the form of carbon isotopic composition of kerogens ~2.7 Ga^4^ and methane-bearing fluid inclusions ~3.46 Ga^5^. Therefore, the time of the onset of microbial methane production and the relative contributions of microbial and abiogenic sources to Archaean atmospheric methane remain uncertain. The case for a microbial methane contribution would be strengthened by molecular clock estimates showing the divergence of methane-producing microbes predates their proposed geochemical signature. Few such studies have been conducted, resulting in a range of dates for the origin of microbial methanogenesis spanning the early Precambrian (e.g., 3.05-4.49 Ga^1^, 3.46-3.49 Ga^6^, ~3.45 Ga^7^, 2.97-3.33 Ga^8^, and a much younger 1.26-1.31 Ga^9^). Estimating divergence times requires calibration points from the geological record^10^: body or trace fossils attributable to a clade’s crown group, preferably by phylogenetic analysis^11^, or (more controversially) preserved traces of organic biomarkers that may be diagnostic for certain clades^12,13^. There is no such direct evidence, however, for Archaea in deep time (let alone methanogens nested deeply within Euryarchaeota). Recent molecular phylogenies suggest eukaryotes may have evolved from within a paraphyletic Archaea^14^. However, the lack of consensus as to the placement of eukaryotes^15^, and the long branch separating eukaryotes from all other groups, make direct fossil calibration of Archaea using crown group eukaryotic fossils problematic. Without geological constraints, confidence in divergence estimates rests entirely on the unconstrained rate models and root priors used, which are sensitive to lineage-specific rate changes^16^, and cannot be internally cross-validated.

In this work, we employ a horizontal gene transfer (HGT) event to date the divergence of methanogens. The HGT was donated from methanogens to the ancestor of Cyanobacteria, microbial clade with the oldest fossils with likely crown-group affinities known in the entire Tree of Life. This extends the use of fossil-calibrated relaxed molecular clocks to archaeal evolution, enabling methods comparable to those validated in studies of metazoan evolution. As calibrations from the rock record are essential for accurate molecular clock inferences, particularly for ancient splits^10^, their inclusion permits more accurate and precise dating of the earliest methanogens.

HGT events represent temporal intrusions between genomes, establishing a cross-cutting relationship determining the relative age of the donor (older) and recipient (younger) clades^17^. Previous work has argued for the relative ages of clades^18–20^, or has used HGT events as secondary calibrations for molecular clock studies^21,22^. Caution must be applied when importing secondary divergence estimates from prior molecular clock studies, as they may propagate errors associated with the original estimate, leading to false precision^23,24^. Furthermore, basing a molecular clock solely on donor-recipient logic fails to incorporate the observed reticulating branch length. This is relevant as it is impossible to ascertain whether the HGT occurred near the divergence of the recipient’s total group, near the diversification of its crown group, or at any time along its stem lineage. As stem lineages can represent very long time intervals for major microbial clades, their omission may dramatically impact date inferences.

Previous phylogenetic analyses have shown that the *smc, scpA*, and *scpB* genes (together encoding proteins that form the SMC complex, required for chromosome condensation in many microbial groups), were transferred to Cyanobacteria from a euryarchaeal donor^25,26^. With improved taxon sampling and accounting for long branch attraction artifacts, we show strong phylogenetic support that SMC complex genes were transferred in a single evolutionary event from a sister lineage of Methanomicrobiales to the ancestor of Cyanobacteria (**Fig. 1, Supplementary Figs. 1-4, Supplementary Table 1**).

**Figure 1.**
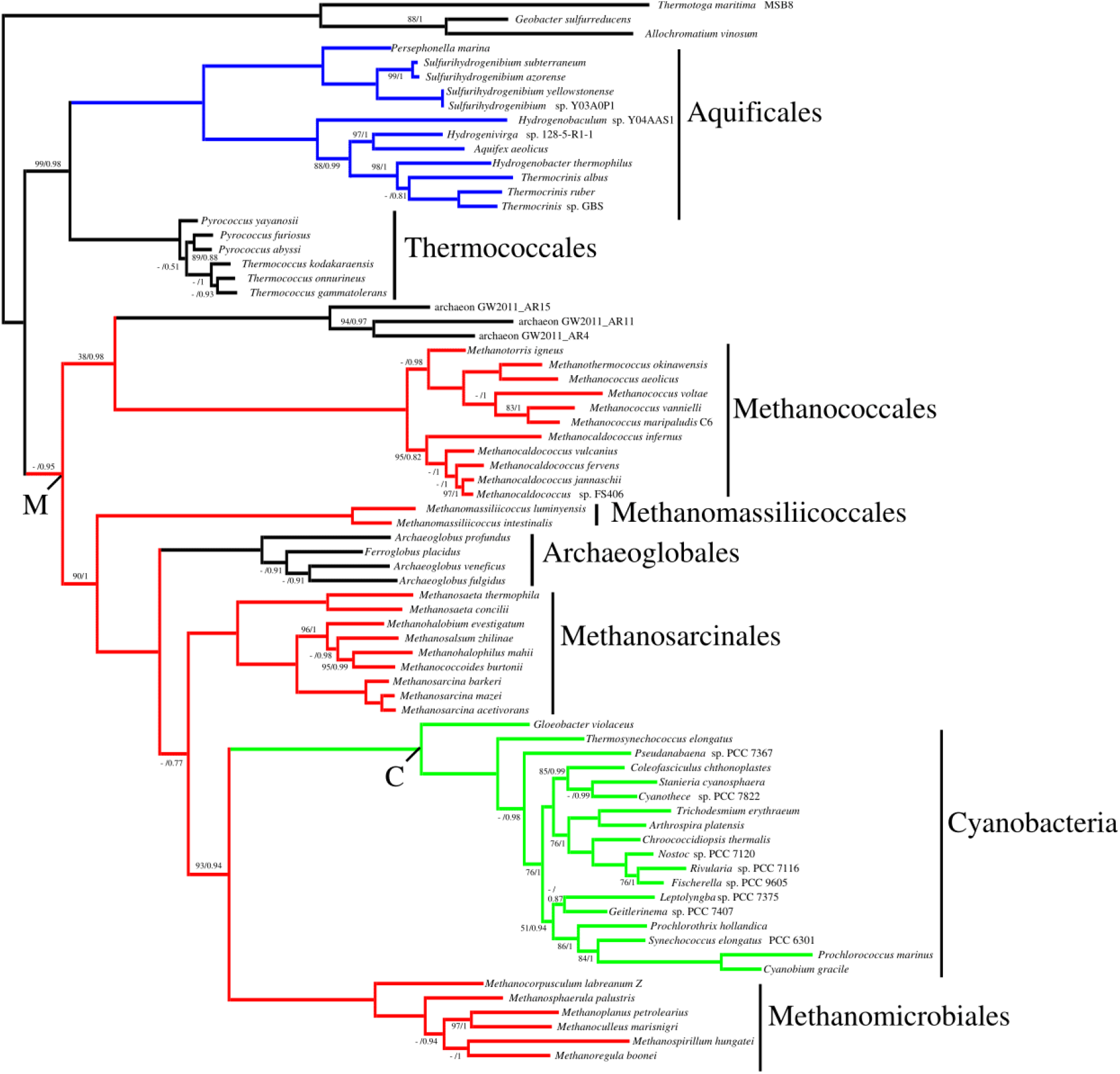
Concatenated PhyloBayes gene tree of *smc, scpA*, and *scpB* for Euryarchaeota (methanogenic lineages in red, node labeled M), with HGT to Aquificales (blue), and Cyanobacteria (green, labeled C). Numbers at nodes represent bootstrap percentages (unlabeled nodes have 100% support) / posterior probabilities (unlabeled nodes have pp = 1.00). Monophyly of the methanogen node was recovered by PhyloBayes with the CAT20 substitution model (pp = 0.95), and was not recovered by RaxML with the LG4M + G model (bootstrap = 52% for alternative topology), hence the Bayesian topology is shown. Nodes not supported in the ML topology are indicated by a dash (-).

## Results

To link the HGT event with the species topology of both methanogens and Cyanobacteria, we concatenated 1) aligned SMC complex sequences for Cyanobacteria and Euryarchaeota with 2) ribosomal protein sequences for Euryarchaeota (expected to reconstruct the Euryarchaeota species tree; listed in **Supplementary Table 2**) and 3) ribosomal protein sequences for Cyanobacteria as three separate partitions in a composite alignment. This composite alignment allows the reticulating branch length for SMC complex evolution to be included in dating analyses, while the topology and branch lengths of the respective euryarchaeal and cyanobacterial clades are inferred from the far more extensive site information within ribosomal datasets. The composite alignment maximally captures the sequence information required to infer divergences of the donor and recipient lineage, as well as sequence information supporting the length and placement of the reticulating branch.

Pairwise distances (**Supplementary Fig. 5**) suggest that the SMC complex genes are evolving slightly (~30%) faster than ribosomal genes, but at about the same rate in all taxa, including the reticulating branch. Thus inclusion of the HGT does not produce clade-specific lineage effects, and the HGT is appropriate for concatenation in a composite alignment. Observed heterotachy may have been imposed by the HGT event itself, which may impact rate estimates along this branch. To test this potential impact, simulated alignments were generated using artificially halved and doubled reticulating branch lengths (**Supplementary Fig. 6**). On average, doubling the reticulating branch length decreased the age of the cyanobacterial crown group by ~77 Myr, and increased the age of the methanogen donor clade by ~87 Myr. Halving the reticulating branch length increased the age of the cyanobacterial crown group by ~72 Myr, and decreased the age of the methanogen donor clade by ~62 Myr. Given the large variances associated with each of these age estimates, impacts on divergence times were relatively small.

Previous studies have shown Bayesian dating approaches may be robust to extensive missing sequence data^27^. We further explored the suitability of composite alignments with large blocks of missing data (where entire clades lack all 30 ribosomal sequences) using simulations (**Supplementary Fig. 7**). The mean age estimates for crown Cyanobacteria were slightly older when missing data were included, increasing the age of crown Cyanobacteria by 2.6%; however, the mean age estimates for the donor clade were not significantly affected. Therefore, missing data have a small impact on age estimates, but this level of significance does not propagate to deeper nodes.

The accuracy of divergence times estimated individually from the Euryarchaeota species tree (uncalibrated relaxed clock only) differed substantially from the SMC gene tree (incorporating the HGT into the analysis) and the composite alignment result (**Fig. 2A**). Based on the species tree alone, methanogens are estimated to have diverged within the Paleoarchaean (mean 3.53 Ga ± SD of 163 Myr, minimum 3.24 Ga). Analyses of the SMC complex alone (mean 3.96 Ga ± 236 Myr, minimum 3.46 Ga) and the composite alignment (mean 3.94 Ga ± 228 Myr, minimum 3.51 Ga) both yield older age estimates for methanogens, in the Eoarchaean. Precision of the latter two analyses is similar for deeper nodes and slightly lower in the composite alignment for Cyanobacteria. Across all calibration sets, the effective prior distributions are similar to the posterior results for the root and methanogen node, but differ slightly for Cyanobacteria, where the posterior is included within a broader effective prior (**Supplementary Table 3** and **Supplementary Fig. 8**). This indicates that prior specification is responsible for age results at most nodes, but the sequence data are also informative for some nodes. In the Euryarchaeota species tree alone, the donor node (Methanosarcinales + Methanomicrobiales) was not significantly older (2.46 Ga ± 158 Myr) than estimates for crown Cyanobacteria from other analyses (mean 2.32 Ga ± 180 Myr; below), which is unlikely as the donor node *must* be older than the recipient’s crown group^22^. This necessary adjustment makes up for the slight decrease in precision when the HGT partition is added, providing an additional calibration and increased accuracy. The advantage of adding ribosomal alignment blocks to the HGT is thus the incorporation of taxa and outgroups that lack the SMC complex genes, allowing us to infer the ages of more ancient nodes, including the divergence of methanogens, and their earliest diversifications.

**Figure 2.**
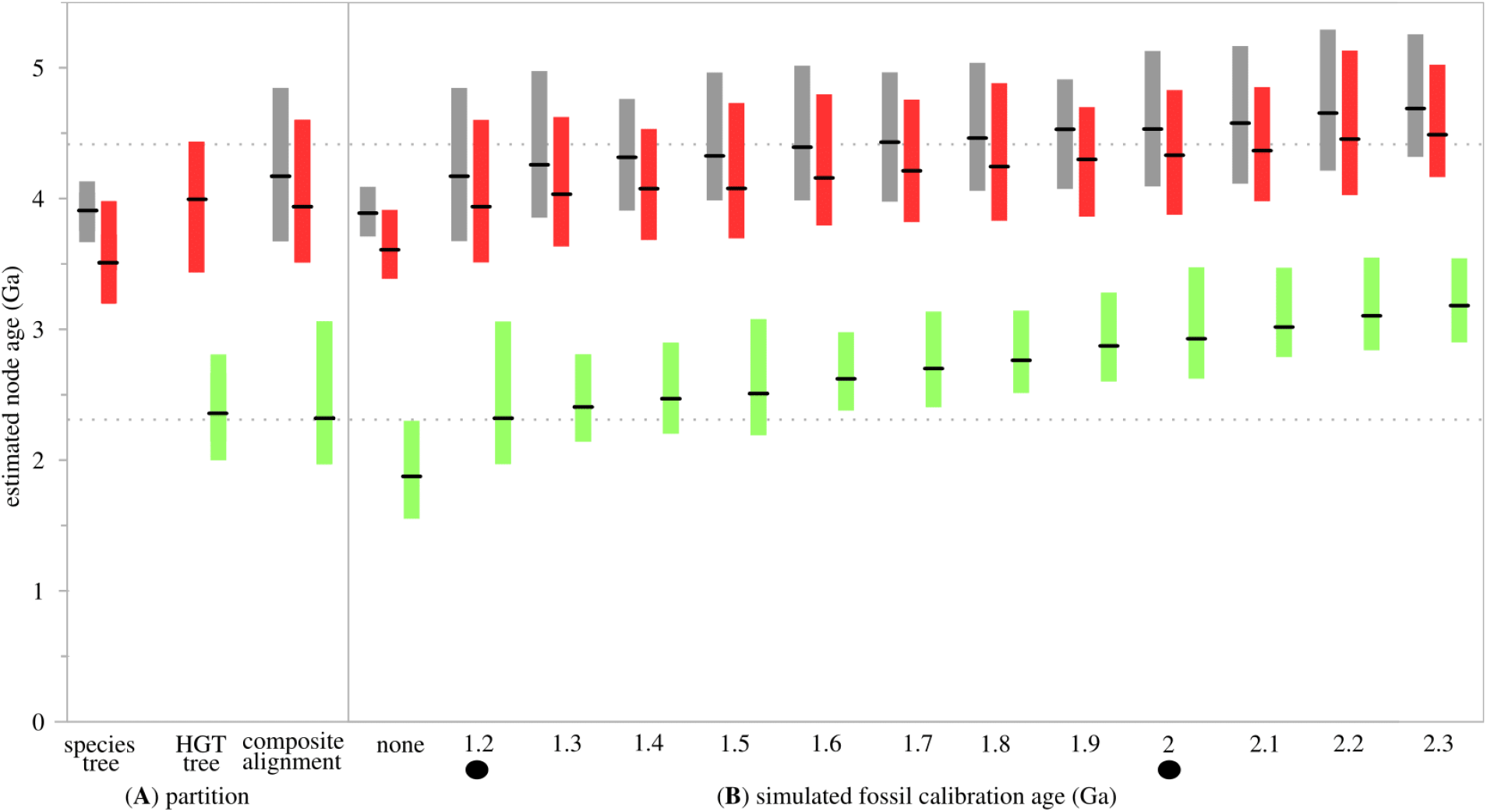
Comparisons of 95% CI date estimates for Cyanobacteria (green; corresponding to node C in **Figs. 1** and **3**), methanogenic Euryarchaeota (red; corresponding to node M in **Figs. 1** and **3**), and crown Euryarchaeota (grey; corresponding to node R in **Fig. 3**) obtained from fixed topologies, with ages reconstructed in PhyloBayes using the CAT20 substitution model, UGM molecular clock model with uniform rates across sites, and gamma distributed root prior of 3.9 Ga with a standard deviation of 230 Myr. Lower dotted horizontal line represents the GOE^29^; upper dotted line represents the oldest zircons^30^. **(A)** Separate effects of the Euryarchaeota species tree (does not include Cyanobacteria), HGT gene tree (does not include a root estimate, because the SMC complex is not found in all methanogens), and composite alignment. The HGT gene tree and composite alignment are calibrated with 1.2 Ga akinete fossils^33^. **(B)** Effect of simulated fossil constraints. Filled circles indicate empirical fossil ages^28^.

## Discussion

Divergence time estimates calibrated by a 2.0 Ga fossil akinete (rod-like resting cell^28^ are extremely old (**Fig. 2B**), with the age of Cyanobacteria (mean 2.93 Ga ± 161 Myr, minimum 2.62 Ga) substantially predating the Great Oxygenation Event (GOE; 2.33 Ga^29^), and with the age of the methanogen ancestor tipping into the Hadean (mean 4.33 Ga ± 240 Myr, minimum 3.88 Ga). In this analysis, the age of Euryarchaeota (mean 4.53 Ga ± 252 Myr, minimum 4.09 Ga) violates the maximum prior applied to the root, estimating a most likely ancestor age older than the oldest zircons (4.38 Ga^30^), and possibly older than the Earth itself. The maximum plausible fossil age for total-group Nostocales (before resulting mean estimates violate the root prior; **Fig. 2B**) corresponds to ~1.7 Ga, a similar age to proposed akinete material from the McArthur Group of Northern Australia^31^. As the validity of the 2.0 Ga microfossil has been questioned^32^, we also calibrated the same node on our tree with a younger 1.2 Ga fossil akinete^33^, which has greater morphological evidence^32^, and is the most conservative estimate discussed more extensively below (**Fig. 3**). This calibration results in age estimates for Cyanobacteria (mean 2.32 Ga ± 180 Myr, minimum 1.97 Ga) very close to and potentially younger than GOE, microbial methanogenesis in the Eoarchaean (mean 3.94 Ga ± 228 Myr, minimum 3.51 Ga), and a correspondingly early age for Euryarchaeota (mean 4.17 Ga ± 228 Myr, minimum 3.67 Ga).

**Figure 3.**
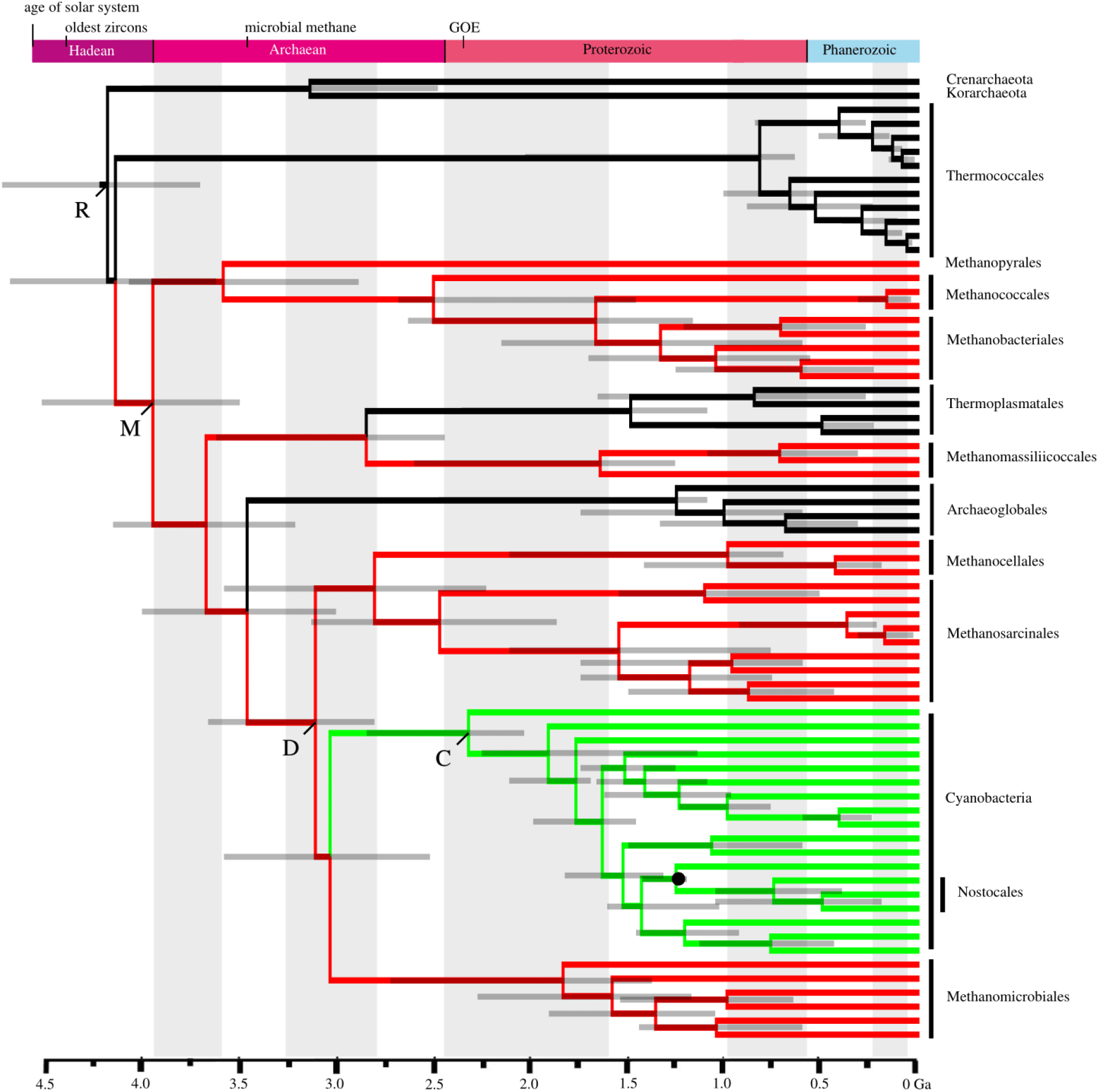
Most conservative divergence time estimates of Euryarchaeota + Cyanobacteria from composite alignment, estimated in PhyloBayes, using the CAT20 substitution model, UGM molecular clock model, and gamma distributed root prior of 3.9 Ga with 230 Myr standard deviation. Branches for Euryarchaeota and Cyanobacteria are determined by ribosomal alignment partitions; the reticulating branch of the cyanobacterial stem is determined by the SMC (HGT) partition. Letter labels on nodes as follows: R = root, M = methanogens, D = HGT donor clade, C = Cyanobacteria. The fossil calibration from 1.2 Ga akinetes^33^ is indicated by a filled circle. Bars on nodes indicate 95% confidence intervals. Note the GOE age is based on new sulfur isotope measurements^29^, and is thus younger than the base of the Proterozoic.

Although only a single fossil calibration was used for this analysis, it may still improve accuracy and precision where among-lineage rate variation is accounted for jointly with the root prior^34^. In simulations, age estimates for Cyanobacteria are substantially more accurate when a fossil calibration is added (**Fig. 2B**), while deeper nodes in Euryarchaeota are less influenced. The 95% confidence intervals calculated from empirical fossils (1.2 and 2.0 Ga) overlap for the ages of Euryarchaeota and microbial methanogenesis, but not for Cyanobacteria, illustrating the importance of sensitivity analysis for clades such as Cyanobacteria with ghost ranges dependent upon phylogenetic interpretation of fossil discoveries^35^ and the use of (relatively) “safe but late” constraints^23^. Note that the GOE itself was not used as a calibration, as different age estimates of Cyanobacteria are contradictory about the relationship between the timing of oxygenic photosynthesis and the age of Cyanobacteria themselves^36^.

Within Euryarchaeota, Methanopyrales, Methanococcales, and Methanobacteriales diverge earlier than Methanomicrobiales, Methanosarcinales, and their relatives (**Fig. 3**). We conservatively estimate the emergence of methanogens as 3.94 Ga ± 228 Myr at the youngest, and the split between Methanosarcinales and Methanomicrobiales (the closest split to the HGT) at 3.10 Ga ± 195 Myr. Therefore, any proposed scenario of a late origin of microbial methanogenesis in the Mesoarchaean through Proterozoic^8,9^ violates the youngest possible calibrated molecular clock estimate (95% CI younger bound of 3.51 Ga), in addition to geochemical evidence^5^. Recently, archaeal clades outside of Euryarchaeota, the uncultured Bathyarchaeota and Verstraetearchaeota, were found to possess genes involved in methane metabolism^37,38^, thus the absolute origin of microbial methanogenesis could be substantially older than Euryarchaeota. Although our analysis is agnostic regarding the ancestral metabolism of Archaea, an older evolutionary history for microbial methanogenesis does not refute the hypothesis of an Archaean microbial methane greenhouse.

A substantial microbial methane greenhouse likely only contributes to Archaean warming of the Earth if 1) methanogenesis evolved early enough, which is consistent with our age estimates, and 2) the divergence time of methanogens predates that of the diversification of poorly characterized microbial taxa involved in anaerobic oxidation of methane (AOM; usually comprising communities of Bacteria and Archaea living together). AOM taxa are of interest because their metabolism can alter the carbon isotope signature of methane produced by microbes^5,39^ in the opposite direction from microbial methanogenesis^5,39^. Furthermore, AOM removes a substantial fraction of methane from sediments, which could effectively erase any geochemical signature of Eorchaean microbial methanogenesis^40,41^. Divergence time estimates for AOM taxa alone could not directly support the existence of microbial methanogenesis, as they can also metabolize abiotic sources of methane^42^. Thus, comprehensive estimates of divergence times for both methanogenic Euryarchaeota and AOM taxa could together constrain the Archaean “methane greenhouse window”, by permitting a narrower, independent interpretation of isotopic data.

As in biostratigraphy, in which index fossils are used to correlate and calibrate rock formations worldwide, a well-supported HGT event from a clade of interest into a fossil-bearing clade permits a direct link between divergence estimates and geological history. Combining data from genes with both reticulate and vertical histories into a single alignment complements other recent developments in calibrating microbial evolution^20,22,43–45^. Our results strongly support the appearance of major methanogen lineages predating the emergence of crown group Cyanobacteria. Our divergence estimates for Euryarchaeota are consistent with previous hypotheses proposing a role for microbial methane in warming the Archaean Earth. With the growing importance of time-calibrated phylogenies in evolutionary inference^46,47^, these methodological developments help to overcome the limitations of the sparse microbial geologic record, and indicate their potential utility in resolving the comparative natural history of microbial clades across the entire Tree of Life.

## Methods

### Data Matrix Construction

The *smc, scpA* and *scpB* proteins form a complex required for chromosome condensation in many microbial groups. Genes encoding these proteins within Cyanobacteria have previously been identified as having been transferred from within Archaea^25,26^. We queried NCBI’s nr database using BLASTp for homologs of *smc, scpA*, and *scpB* proteins in each member of Euryarchaeota with a sequenced genome (except the species-rich Halobacteriales, for which we selected eight representatives), and representative Cyanobacteria from all orders. Previously reported SMC homologs within Aquificales likely representing an additional HGT from Thermococcales^26^ were also included. No *scpB* sequences were found in Aquificales or Halobacteriales. Protein sequences for each homolog were individually aligned in MUSCLE v3.7^48^. The *smc* protein contained two large poorly aligned regions, representing coiled-coil domains^25,49^. These regions were removed via alignment masking using GUIDANCE^50^, leaving 729 aligned sites. For the two *scp* proteins (which are much shorter, with limited phylogenetic informativeness; **Supplementary Table 4**), we elected not to mask poorly aligned regions in light of recent work indicating trees resulting from this process may be of decreased quality^51^.

### Phylogenetic Analysis

Individual gene trees were constructed with RaxML v1.8.9^52^ using the LG4M + G substitution model^53^. All three HGT genes were concatenated with FASconCAT v1.0^54^, analyzed in RaxML with 100 bootstrap replicates, and in PhyloBayes v3.3f using two chains and the CAT20 site-dependent model^55,56^. The CAT20 model was used because preliminary analyses using the full CAT model did not reach convergence. An automatic stopping rule was implemented, with tests of convergence every 100 cycles, until the default criteria of effective sizes and parameter discrepancies between chains were met (50 and 0.3, respectively). Trees and posterior probability support values were then generated from completed chains after the initial 20% of sampled generations were discarded as burn-in.

### Composite Alignment

A composite alignment was constructed to date the origin of methanogens, by concatenating 1) aligned SMC complex sequences for Cyanobacteria and Euryarchaeota (1,778 amino acids) with 2) ribosomal sequences for Euryarchaeota (adding representatives of clades without identified SMC homologs, i.e. Methanobacteriales, Methanocellales, Methanopyrales, and Thermoplasmatales) and 3) ribosomal sequences for Cyanobacteria, as three separate partitions (14,366 amino acids total). Specifically, 30 ribosomal proteins (**Supplementary Table 2**) were identified by BLASTp, aligned separately in MUSCLE, then concatenated. Separate partitions for cyanobacterial and archaeal ribosomal proteins are used to provide more informative sites for estimating evolutionary relationships and rates in these groups, without introducing phylogenetic conflict with the HGT partition. Using this approach, only SMC complex sequences determine cyanobacterial placement ‘within’ Euryarchaeota along the reticulating HGT branch. Note that SMC sequences from Aquificales were omitted from these analyses, as this additional putative HGT event is uninformative in this investigation. The concatenated topology was estimated with RaxML using the LG4M + G model and PhyloBayes using CAT20.

### Fossil Calibration

To produce a divergence estimate, we applied a time constraint within Cyanobacteria, derived from fossil resting cells (akinetes; genus *Archaeoellipsoides*) similar to the cyanobacterial clades Nostocales (morphological subsection IV) and Stigonematales (subsection V), from the 2.0 Ga Franceville Group of Gabon^28,31,57,58^. There are too few morphological characters to determine a crown-group position of this fossil^32^, so we assigned the fossil minimum age to total-group Nostocales (i.e. the clade in our tree including Nostocales and Stigonematales^32^, and their sister group Chroococcidiopsidales). As the affinities of Paleoproterozoic *Archaeoellipsoides* have been questioned^32,59^, we also tested a less controversial younger fossil of the same genus with a more similar size to members of total-group Nostocales, from the 1.2 Ga Dismal Lakes Group of Northwest Canada^33,59^. To measure the effect of using different *Archaeoellipsoides* fossil ages on divergence time estimates, we simulated calibrations for total-group Nostocales at 100 Myr intervals between 1.3 and 2.3 Ga (in addition to the empirical fossil dates at 1.2 and 2.0 Ga). Unlike some previous analyses^57,58^, we did not include the age of the GOE as a calibration on the age of Cyanobacteria. Our approach permits an estimate of the age of Cyanobacteria independent of the onset of atmospheric oxygenation^36^.

### Divergence Time Estimation

Divergence times were estimated in PhyloBayes using a fixed topology from the RaxML composite alignment result, the CAT20 substitution model, and the uncorrelated gamma multipliers (UGM) relaxed clock model^16^. The UGM model allows substitution rates to vary across the tree, and assumes there is no autocorrelation of evolutionary rates across deep branches^16^. Therefore, this model is suited to modeling rate changes associated with HGT events along a reticulating branch. Rates across sites followed a uniform distribution, and the prior on divergence times was uniform.

The root was calibrated with a gamma distributed prior with a mean of 3.9 Ga and SD 230 Myr (range from 4.36 to 3.44 Ga); this constraint was calculated as the mean of the maximum root age of 4.38 Ga (oldest zircons, approximating the age of habitable Earth^30^) and minimum of 3.46 Ga (oldest traces of microbial methane^5^). We selected a gamma distribution rather than uniform, because we do not assume it would be equally likely that the last common ancestor of Euryarchaeota diverged at either the maximum or minimum age (i.e. the tails are less likely). The superiority of “soft” calibration densities has been discussed previously^60^. It is not circular to use microbial methane traces as a younger bound on the root, because this constraint only presupposes the methane traces are 1) archaeal and 2) biogenic, and does not specifically constrain the age of any clade within Archaea, including methanogens (i.e., the ancestor of known methanogens may be either younger or older than 3.46 Ga, as the prior is not directly placed on its node). Each fossil age (above) was used as a hard-bound minimum constraint on a uniform age prior, which is appropriate (despite soft bounds on the root, above) due to the extreme antiquity and limited character information from these calibrations. Other validatory analyses, including varying the molecular clock model resulted in minimal changes to divergence time estimates. Comparisons of estimated CIs to the effective prior^61^ were also made by removing sequence data using the -prior flag in PhyloBayes (**Supplementary Fig. 8** and **Supplementary Table 3**).

### Data Availability

Supplementary data files are available at Dryad (provisional link: http://datadryad.org/review?doi=doi:10.5061/dryad.m371v).

## Acknowledgments

We thank D. Pisani and M. dos Reis for improving the manuscript with their helpful comments, D. Gruen, C. Magnabosco, D. Rothman, and B. Schirrmeister for discussions, and G. Shomo for assistance with the Engaging Cluster at MGHPCC. We acknowledge support from Simons Foundation Collaboration on the Origin of Life #339603 to G.P.F. and NSF EAR-1615426 to G.P.F. and J.M.W.

## Author Contributions

J.M.W. and G.P.F. designed research and performed data analysis. J.M.W. drafted the manuscript with assistance from G.P.F.

## SUPPLEMENTARY INFORMATION

### Long Branch Attraction Sensitivity Analyses

Halobacteriales have an established strong compositional bias, which may result in long branch attraction^1–3^. Omitting Halobacteriales from the concatenated SMC complex alignment (**Supplementary Table 1**) significantly improved phylogenetic support that the transfer of SMC proteins to the ancestor of Cyanobacteria was most likely from a sister lineage of Methanomicrobiales (**Fig. 1**).

### Pairwise Distances

Pairwise distances between all shared taxa were extracted from the phylogenies generated from the SMC complex (HGT) and composite alignments (**Fig. 1** and topology of **Fig. 3**, respectively) using T-REX^4^. A plot of these distances (**Supplementary Fig. 5**) shows a generally linear relationship between alignments for methanogen and cyanobacterial groups, with consistently slightly longer branches within the SMC complex gene tree.

### HGT Branch Length Simulations

We tested whether the branch length of the HGT itself had a significant effect on the assessed divergence times. Ten simulations were generated for each of two trees, with the same topology as the composite alignment (i.e., that depicted in **Fig. 3**), but in which the reticulating branch length was altered, by either doubling or halving its length. In this way, the effect of rate changes along a reticulating branch induced via HGT on the molecular clock model could be observed. For each branch length simulation, a divergence time analysis was performed on the simulated sequences in PhyloBayes^5^, using the same parameters as the empirical data and the fossil akinete calibration from 1.2 Ga^6^. All runs on simulated data converged.

### Missing Data Simulations

To test the role of missing data in the composite alignment, a control simulation was created. Ten simulated alignments were generated using the PAML4 module evolver^7^, using the LG model with eight discrete gamma-distributed categories and a shape parameter of alpha=0.69, as best fit to the concatenated ribosomal-SMC protein composite alignment by ProtTest^8^. Amino acid frequencies were matched to those observed within the composite alignment. Sequences were simulated along the inferred maximum likelihood tree recovered from the concatenated dataset (i.e. the topology in **Fig. 3**). Sequences were initially simulated for 11223 sites, to match the full number of sites within the composite alignment that have less than 50% gaps within sequence blocks.

For each simulated alignment, blocks of of equal length and taxonomic distribution to the regions of missing data within the observed alignment (e.g. the blocks between euryarchaeal and cyanobacterial ribosomal proteins) within composite alignments were replaced with gaps. 6060 sites in the Cyanobacteria ribosomal partition were replaced with gaps, corresponding to euryarchaeal ribosomal sites absent in Cyanobacteria; 575 sites in the Euryarchaeota ribosomal partition were replaced with gaps corresponding to euryarchaeal taxa that do not contain an included SMC homolog; and 4588 sites in the Euryarchaeota ribosomal partition were replaced with gaps corresponding to cyanobacterial ribosomal sites absent in Euryarchaeota. For each control and missing simulation, a divergence time analysis was performed on the simulated sequences in PhyloBayes, using the same parameters and the 1.2 Ga fossil akinete calibration^6^.

**Supplementary Fig. 1.**
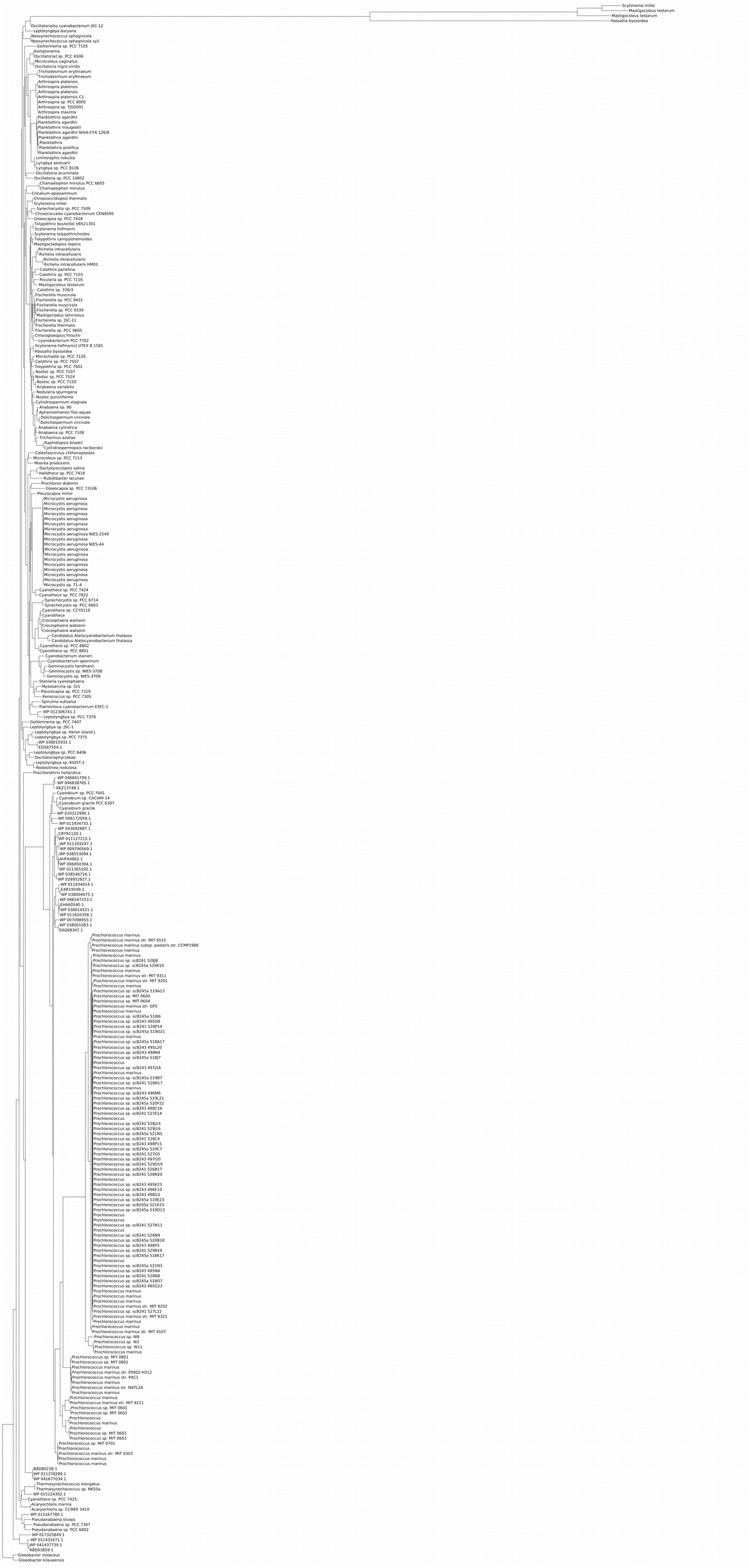
RaxML gene tree of *smc* for all 307 Cyanobacteria taxa available in GenBank, excluding sequences transferred from other bacterial clades.

**Supplementary Fig. 2.**
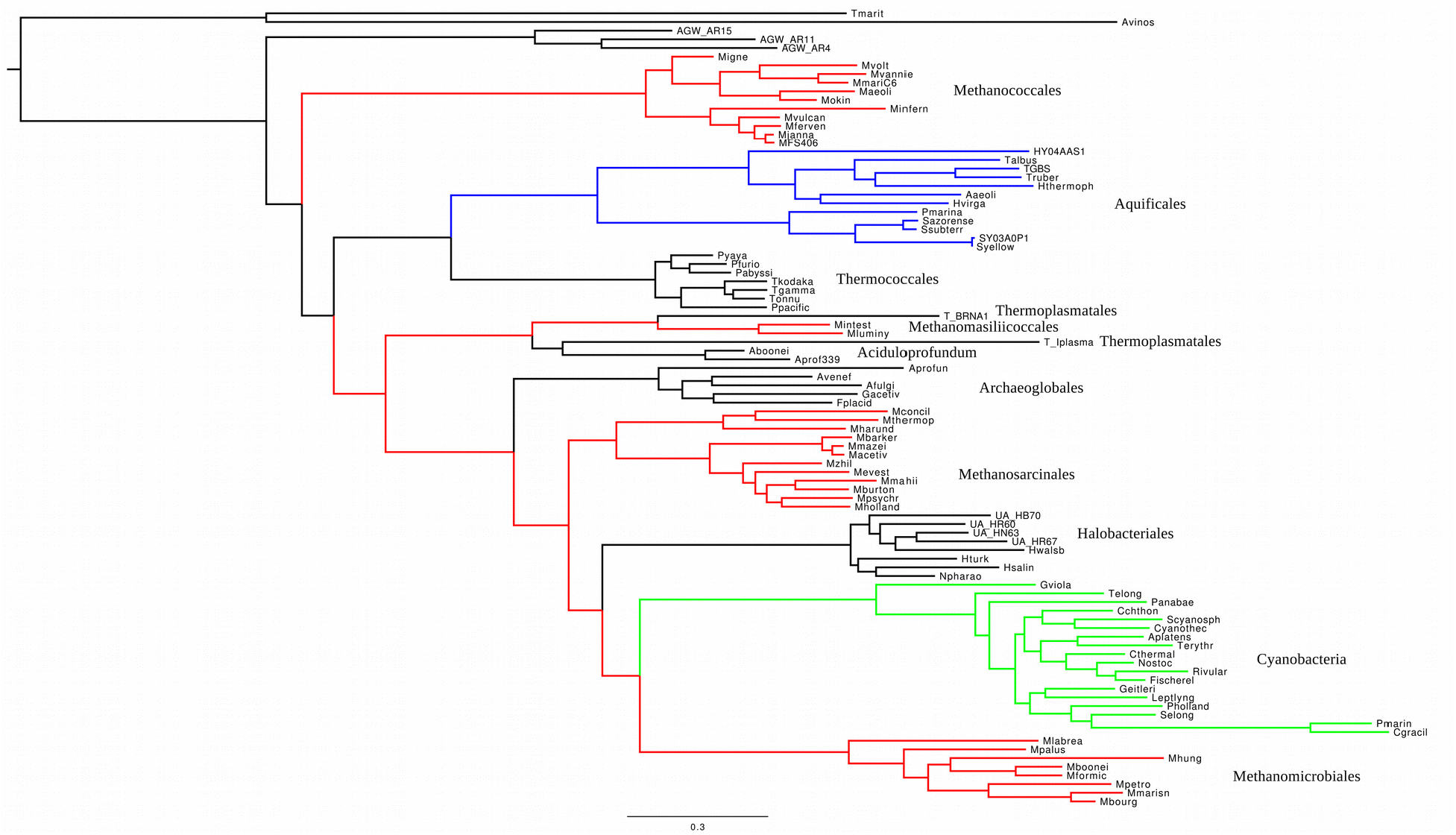
RaxML gene tree of *smc* for Euryarchaeota (methanogenic lineages in red), with HGT to Aquificales (blue), and Cyanobacteria (green).

**Supplementary Fig. 3.**
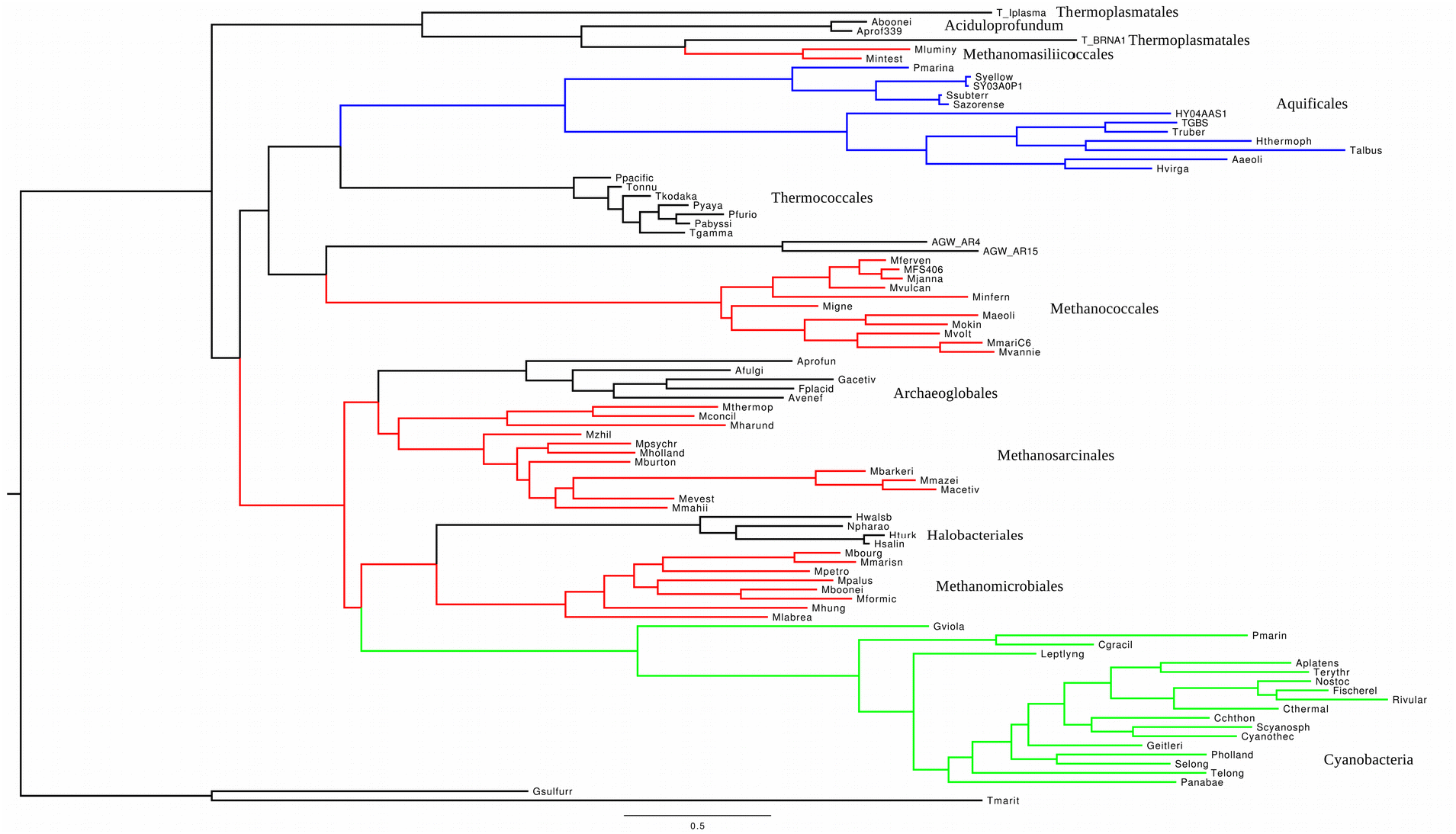
RaxML gene tree of *scpA* for Euryarchaeota (methanogenic lineages in red), with HGT to Aquificales (blue), and Cyanobacteria (green).

**Supplementary Fig. 4.**
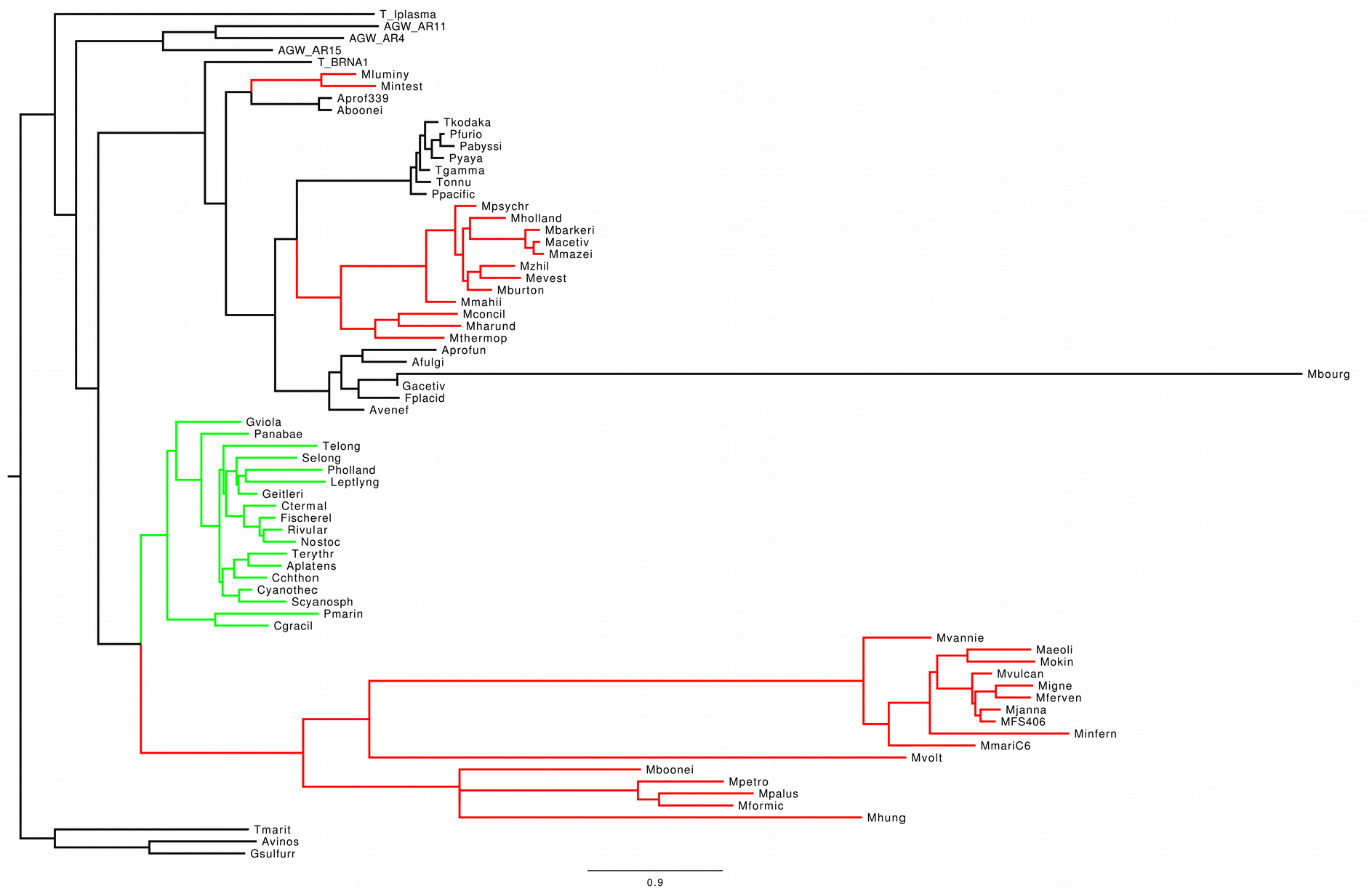
RaxML gene tree of *scpB* for Euryarchaeota (methanogenic lineages in red), with HGT to Cyanobacteria (green).

**Supplementary Table 1.** Bootstrap bipartitions for the concatenated alignment of *smc + scpA + scpB*, calculated in RaxML.

| Cyanobacteria sister group | Bootstrap % | Bootstrap % with Halobacteriales removed |
| --- | --- | --- |
| Methanomicrobiales | 33 | 93 |
| Methanomicrobiales + Halobacteriales | 20 | 0 |
| Halobacteriales | 17 | 0 |
| Methanosarcinales + Halobacteriales | 15 | 0 |
| Methanosarcinales + Halobacteriales + Archaeoglobales | 7 | 0 |
| Methanomicrobiales + Methanosarcinales + Halobacteriales | 4 | 0 |
| Methanomicrobiales + Methanosarcinales + Halobacteriales + Archaeoglobales | 2 | 0 |
| Archaeoglobales | 2 | 0 |
| Methanosarcinales | 0 | 6 |
| Methanomicrobiales + Methanosarcinales | 0 | 1 |

**Supplementary Table 2.** List of ribosomal proteins. These were aligned and concatenated for Euryarchaeota, and separately for Cyanobacteria, to build the composite alignment.

| Large subunit (50S) | Small subunit (30S) |
| --- | --- |
| L1 | S2 |
| L2 | S3 |
| L3 | S4 |
| L4 | S5 |
| L5 | S7 |
| L6 | S8 |
| L10 | S9 |
| L13 | S10 |
| L14 | S11 |
| L15 | S12 |
| L18 | S13 |
| L22 | S14 |
| L23 | S15 |
| L24 | S17 |
| L29 | S19 |

**Supplementary Fig. 5.**
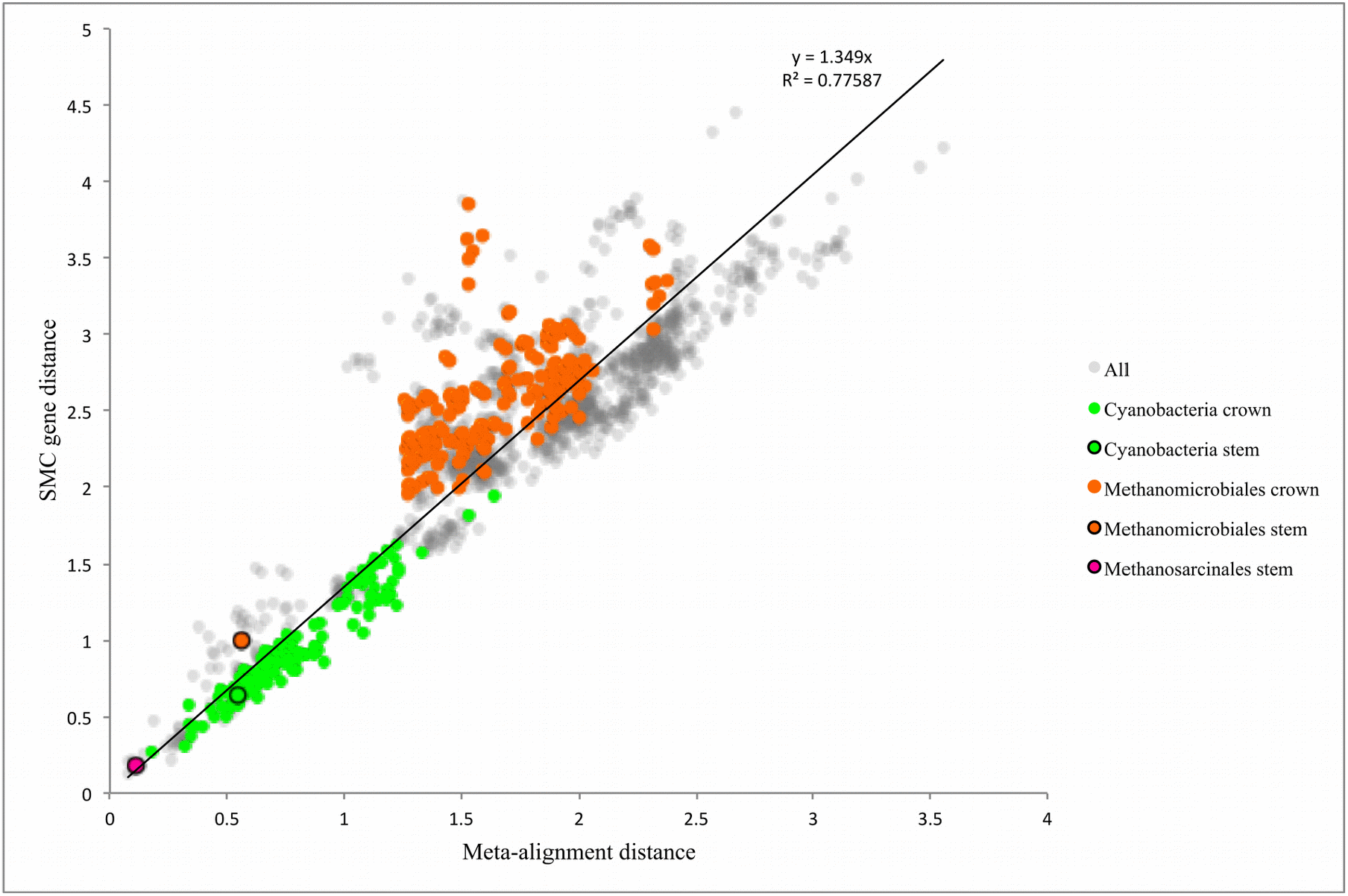
Pairwise distances between branches of the SMC complex gene tree and the composite alignment concatenated tree. Both the root of methanogens and cyanobacterial recipient (green) clades show the same trend, excepting Methanomicrobiales (orange), which are on a disproportionately long branch within the SMC tree. The stem lineage branch lengths are also plotted for Cyanobacteria, Methanomicrobiales, and Methanosarcinales, showing that the long cyanobacterial stem and short Methanosarcinales (pink) stem are proportional between trees and fall on this diagonal (thus no lineage effects are observed among these clades), while the Methanomicrobiales (donor lineage) stem does not (thus this clade may have a lineage specific rate).

**Supplementary Fig. 6.**
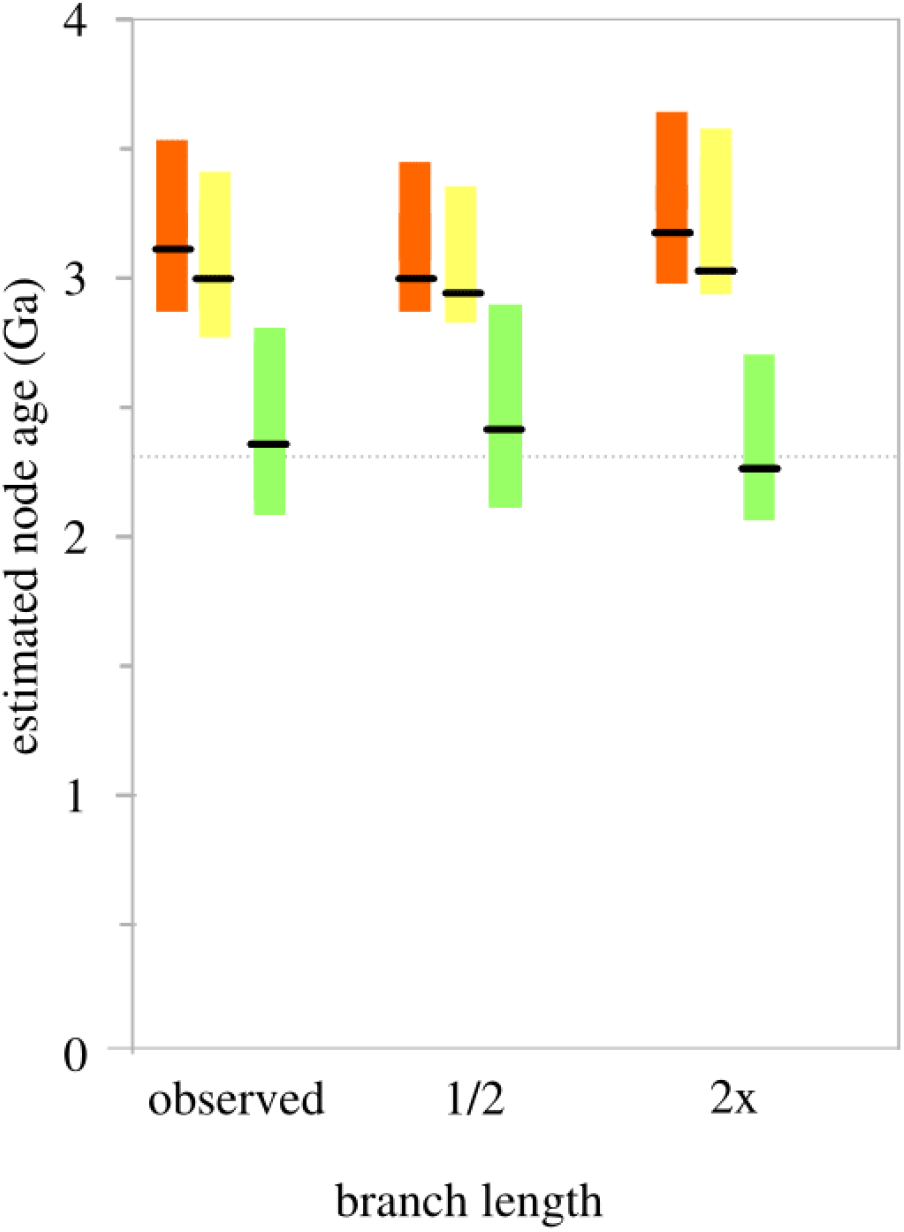
Comparisons of simulated 95% CI date estimates for Cyanobacteria (green), the reticulating node (yellow), and the methanogen donor lineage (orange). Simulations are depicted with the full observed reticulating branch length, with the reticulating branch length half of that observed empirically, and with the reticulating branch length double that observed. Only the mean of 10 simulations is depicted for each mean age and 95% CI.

**Supplementary Fig. 7.**
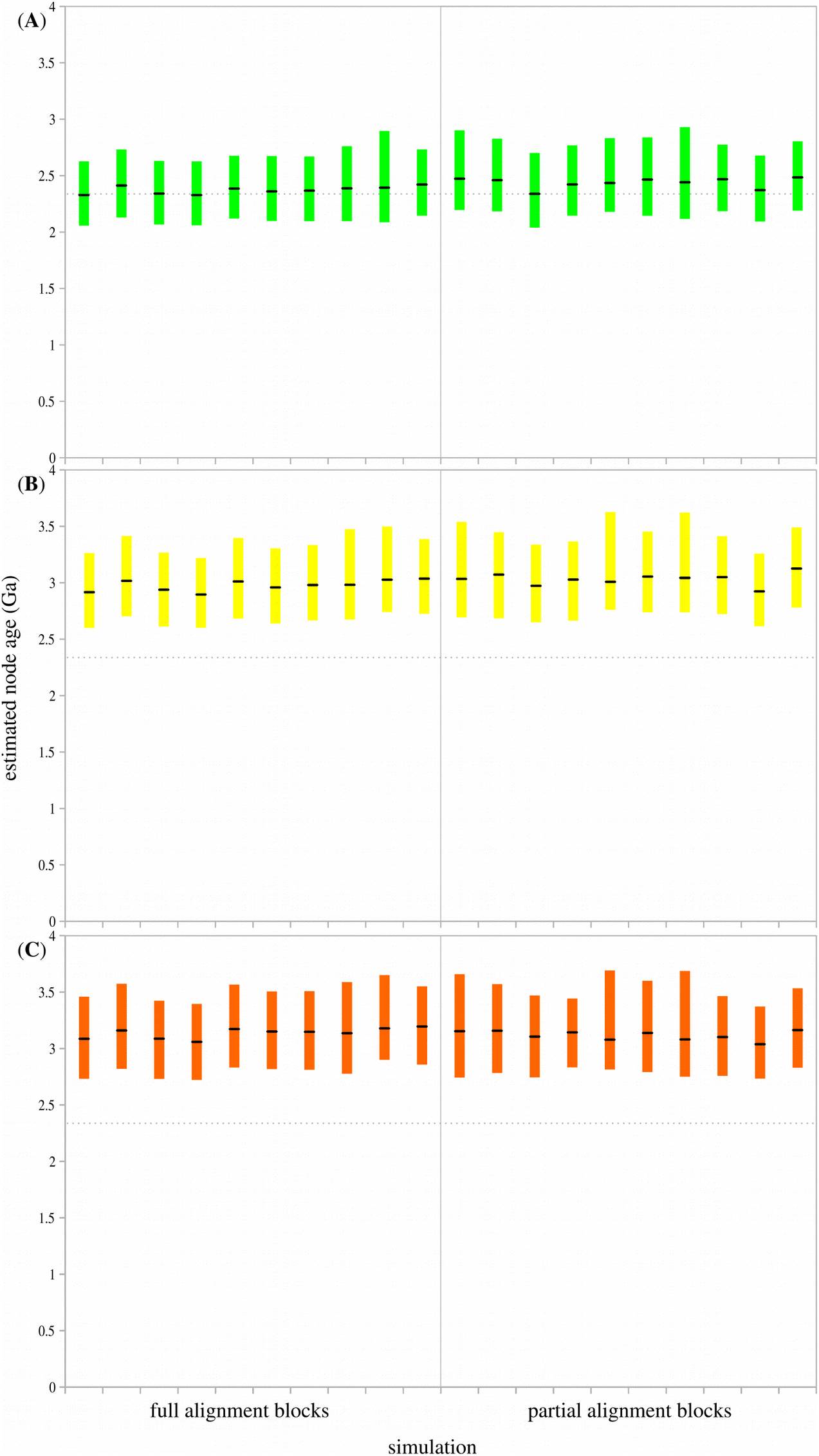
Comparisons of 95% CI date estimates for simulated full length alignments and with blocks of missing data to represent that observed in the composite alignment. All 10 simulations of each dataset depicted have empirical branch lengths from the composite alignment. **(A)** Cyanobacteria (green) simulated age. **(B)** The reticulating node (yellow) simulated age. **(C)** Methanomicrobiales (orange) simulated age.

**Supplementary Table 3.**
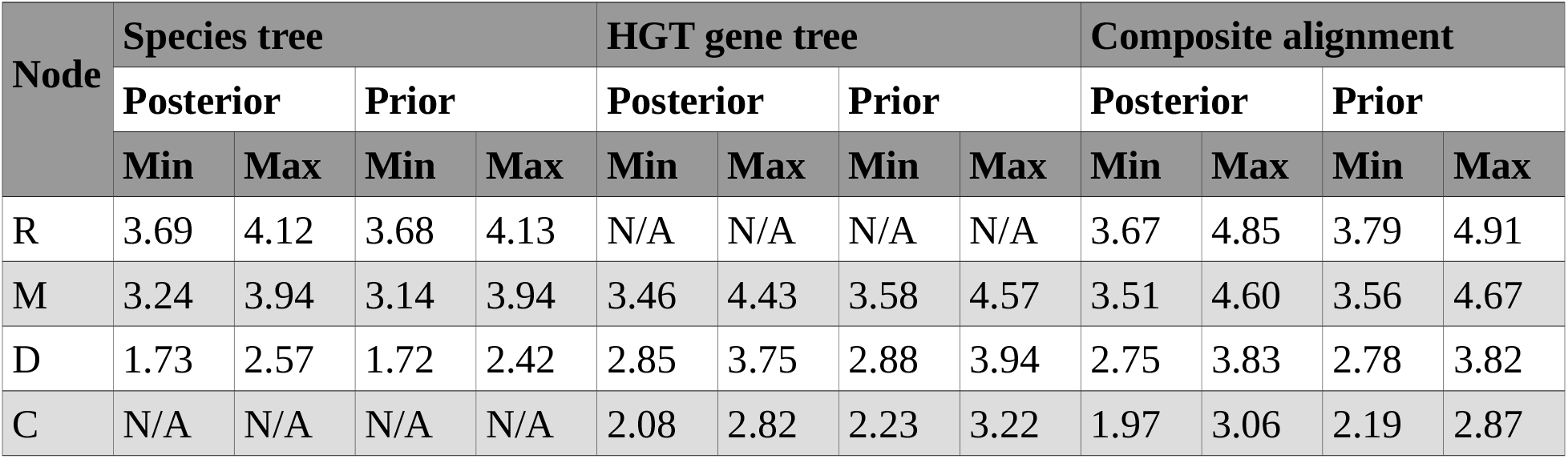
95% posterior CI age estimates in Ga for selected nodes (indicated in **Figs. 1** and **3**), using the same alignments and topologies from **Fig. 2A**. Each estimate was obtained from fixed topologies reconstructed in PhyloBayes, using the CAT20 substitution model, and gamma distributed root prior of 3.9 Ga ± 230 Myr. The ‘Prior’ columns represent estimates resulting from running PhyloBayes with no data (i.e. using the -prior flag). Note the species tree column does not include estimates for Cyanobacteria, because they are not part of the species tree; and the HGT gene tree column does not include estimates of the root age, because the SMC complex is not found in all methanogens.

| Node | Species tree |  |  |  | HGT gene tree |  |  |  | Composite alignment |  |  |  |
| --- | --- | --- | --- | --- | --- | --- | --- | --- | --- | --- | --- | --- |
|  | Posterior |  | Prior |  | Posterior |  | Prior |  | Posterior |  | Prior |  |
|  | Min | Max | Min | Max | Min | Max | Min | Max | Min | Max | Min | Max |
| R | 3.69 | 4.12 | 3.68 | 4.13 | N/A | N/A | N/A | N/A | 3.67 | 4.85 | 3.79 | 4.91 |
| M | 3.24 | 3.94 | 3.14 | 3.94 | 3.46 | 4.43 | 3.58 | 4.57 | 3.51 | 4.60 | 3.56 | 4.67 |
| D | 1.73 | 2.57 | 1.72 | 2.42 | 2.85 | 3.75 | 2.88 | 3.94 | 2.75 | 3.83 | 2.78 | 3.82 |
| C | N/A | N/A | N/A | N/A | 2.08 | 2.82 | 2.23 | 3.22 | 1.97 | 3.06 | 2.19 | 2.87 |

**Supplementary Fig. 8.**
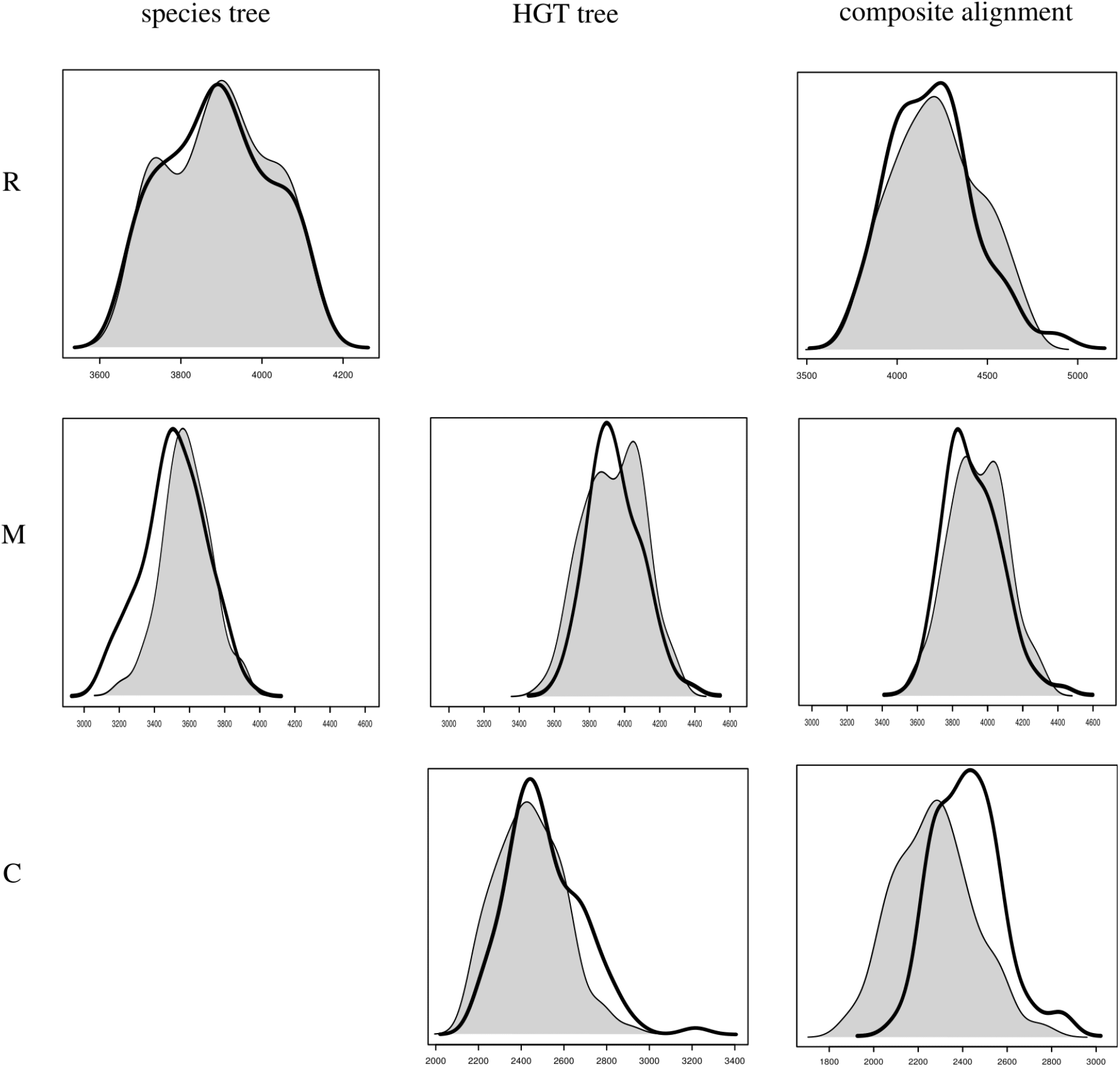
Comparison of posterior probability distributions for divergence times assessed as in **Fig. 2A** (posterior), and using the same analyses under the effective prior (removing sequence data). The posterior analyses are shaded grey; effective priors are superimposed on the same axes with a heavy black line. As in **Fig. 2A**, separate effects of the Euryarchaeota species tree (does not include Cyanobacteria), HGT gene tree (does not include a root estimate, because the SMC complex is not found in all methanogens), and composite alignment. Letter labels refer to the following nodes: R = root, M = methanogens, C = Cyanobacteria.

**Supplementary Table 4.** Number of amino acid sites in each single-gene alignment before and after masking with GUIDANCE. Number of sites found in >4 taxa in parentheses.

| Gene | Unmasked | Masked |
| --- | --- | --- |
| *smc* | 1576 (1323) | 729 (588) |
| *scpA* | 560 (423) | 136 (96) |
| *scpB* | 489 (408) | 100 (0) |

